# Resorbable Poly-(D,L)-Lactide Anchorage of Nanoparticulate Mineralized Collagen Materials Maximizes *In Vivo* Skull Regeneration

**DOI:** 10.1101/2025.07.06.663409

**Authors:** Meiwand Bedar, Xiaoyan Ren, Wei Chen, Youngnam Kang, Shahrzad Moghadam, Kelly X. Huang, Kaavian Shariati, Brendan A.C. Harley, Justine C. Lee

## Abstract

**Background:** A clinical demand exists for bone biomaterials that mirror tissue-specific extracellular matrix (ECM) properties and regulate progenitor cell fate. Nanoparticulate mineralized collagen glycosaminoglycan (MC-GAG) scaffolds have demonstrated to promote skull regeneration *in vivo* without addition of growth factors or exogenous progenitor cells, offering a materials-only solution for cranial defect reconstructions. Further enhancement of the safety and regenerative potential of MC-GAG is however necessary for clinical translation. Here, we investigated the regenerative effect of MC-GAG scaffold fixation with a resorbable poly(D,L-lactide) (PDLLA) implant for cerebral protection.

**Methods:** Fourteen-millimeter cranial defects were created in New Zealand white rabbits, divided into four groups: 1) defect only, 2) defect with PDLLA, 3) MC-GAG, and 4) MC-GAG with PDLLA. Initial bone healing assessment was conducted at 3 and 9 months using microcomputed tomography (microCT), histology, reference point indentation, and scanning electron microscopy. Long-term effects were evaluated through *in vivo* microCT imaging at 3, 6, and 9 months, with biomechanical testing of explanted skulls at 9 months.

**Results:** At 3 months, MC-GAG significantly enhanced mineralization compared to empty and PDLLA-treated defects, with even greater mineralization observed for MC-GAG-PDLLA. Histologically, MC-GAG demonstrated trabecular mineralization, while MC-GAG-PDLLA had a structural organization resembling native bone. Biomechanical assessment showed enhanced toughness and stiffness for both MC-GAG and MC-GAG-PDLLA. At 6 months, MC-GAG-PDLLA displayed the highest bone mass, and at 9 months, both MC-GAG-containing scaffolds demonstrated superior bone regeneration compared to controls. By 9 months, PDLLA implants were completely resorbed and MC-GAG-PDLLA exhibited mechanical properties surpassing those of all other groups.

**Conclusions:** Fixation of MC-GAG scaffolds with PDLLA implants improved mineralization and bone regeneration compared to either material alone for cranial defect reconstruction.

**Clinical Relevance Statement:** Cranioplasties are essential for cerebral protection and neurological restoration, but current materials have limitations, necessitating alternative bone biomaterials for improved outcomes.

## Introduction

Cranial defects arising from congenital anomalies, neurosurgical interventions, tumors, trauma, and infections, often necessitate cranioplasties. These procedures are crucial for restoring neurological function, ensuring cerebral protection, and improving psychosocial well-being.(1,2) The current available reconstructive strategies are however rife with limitations. Autologous bone grafting is constrained by limited availability and donor site morbidity and alloplastic materials present an alternative option, albeit with high complication and explantation rates.(1) Consequently, there is an unmet clinical need for regenerative bone biomaterials capable of promoting cranial defect reconstructions in a manner that recapitulates the structure and function of native bone tissue.

While numerous studies have explored the combination of stem cells, growth factors, and scaffolding materials to enhance bone regeneration, none of these strategies have yet translated into clinical practice due to their high costs, long processing times, and unintended side effects.(3, 4) An innovative solution lies in extracellular matrix(ECM)-inspired biomaterials, which harness the instructive capabilities of the ECM to stimulate the regeneration of native bone. This emerging strategy holds promise as a safe and viable option for cranial defect reconstruction.(5)

Our group previously demonstrated the effectiveness of a nanoparticulate mineralized collagen glycosaminoglycan (MC-GAG) material as a cell-free and growth factor-free strategy for stimulating regeneration of rabbit calvarial bone defects.(6–12) Moving forward, further refinement of the material is essential to enhance both safety and regenerative capacity in preparation for clinical translation. A pertinent challenge faced in the clinical implementation is the protection of underlying structures while allowing adequate intracranial pressure to maintain neurological functioning during early bone healing. To address this challenge, the current study investigates how the regenerative capacity of MC-GAG may be influenced by its fixation and overlay with a clinically available resorbable poly-D,L-lactide (PDLLA) implant intended for early-stage cerebral protection.(13–15)

## Materials and Methods

### Fabrication and chemical crosslinking of non-mineralized and mineralized collagen scaffolds

Nanoparticulate mineralized glycosaminoglycan (MC-GAG) scaffolds were fabricated through lyophilization using our previously described techniques (16–18). Briefly, a mixture of microfibrillar type I collagen (Collagen Matrix, Oakland, NJ) and chondroitin-6-sulfate (Sigma-Aldrich, St. Louis, MO) was combined in a phosphoric acid solution containing calcium salts (calcium nitrate hydrate: Ca(NO_3_)2·4H2O; calcium hydroxide: Ca(OH)2, Sigma-Aldrich). The solution was gradually frozen at a rate of 1 °C/min using a freeze dryer (Genesis, VirTis, Gardiner, NY), transitioning from room temperature to −10 °C. After sublimation, the scaffolds underwent sterilization with ethylene oxide and were stored at −20 °C. Scaffolds were cut into 14 mm disks and rehydrated with ethanol, followed by phosphate-buffered saline (PBS) overnight. Subsequently, the scaffolds underwent four hours of crosslinking using 1-ethyl-3-(3-dimethylaminopropyl) carbodiimide (EDAC, Sigma-Aldrich) and N-hydroxysuccinimide (NHS, Sigma-Aldrich), following established protocols (19). To eliminate residual chemicals, the scaffolds were washed and incubated in PBS.

### In vivo rabbit cranial defect reconstruction

Animal experiments were performed in compliance with the USDA Animal Welfare Act and PHS Policy for the Humane Care and Use of Laboratory Animals. Preoperatively, New Zealand White rabbits (2-3 months old; Charles River Laboratories, Wilmington, MA) were given an intravenous injection of 2-5 mg/kg alfaxalone or 0.01 mg/kg dexmedetomidine for induction. The head of each rabbit was shaved and disinfected with Betadine. Anesthesia was maintained with isoflurane gas (1.5–3%) during the procedure and pain control was performed with a subcutaneous injection of 0.02 mg/kg buprenorphine.

Rabbits were divided into 4 groups: 1) defect without reconstruction, 2) defect with PDLLA implant, 3) MC-GAG scaffold only, and 4) MC-GAG with PDLLA implant. The cranial surface was exposed by a midline incision and the overlying parietal periosteum was dissected off of the calvarium. For each rabbit, a 14-mm biparietal full thickness, extradural defect was created by a hand powered trephine and the bone was lifted away without injury to the dura (20). One scaffold was implanted for each rabbit. In groups 2 and 4, 22-mm resorbable Poly(DL-lactide) implants (Resorb x, KLS-Martin L.P., Jacksonville, FL) were fixed with two 3-mm resorbable PDLLA pins (SonicPin Rx, K KLS-Martin L.P.) using SonicWeld Rx advanced ultrasound technology (KLS-Martin L.P.). The incision was closed with 5-0 resorbable Vicryl sutures. Postoperative, rabbits were given 0.03 mg/kg buprenorphine twice-daily for two days, 0.3 mg/kg meloxicam daily for 5 days, and 10 mg/kg orbifloxacin for 10 days.

Three and nine months after implantation, the rabbits were euthanized by intravenous injection of 1 mL of pentobarbital solution (Fatal-Plus®, 390 mg/ml) intravenously via the marginal ear vein. The previous incision was then reopened and the calvarium was exposed. The calvarium including the cranial defect was analyzed grossly and then explanted for micro-computed tomography, histologic, and biomechanical analyses.

### Micro–Computed Tomographic Imaging

Ex-vivo rabbit skull micro-computed tomographic imaging (µCT) was perfomed using the Scanco µCT 35 (Scanco Medical AG, Bruttisellen, Switzerland). Skulls were removed 3 and 9 months after the implantation of scaffolds, fixed in 10% formalin for 24 hours, and then stored in 70% ethanol at 4 °C until scanned. Scans were performed in 70% ethanol using high resolution settings with a source voltage of 45 E (kVp), Intensity (µA) of 177, and a voxel size of 15 µm. Skull areas were contoured to establish volumes of interest and an optimum arbitrary threshold value of 415 was used uniformly for all specimens to quantify mineralized bone areas from surrounding unmineralized scaffold. Three-dimensional reconstruction pictures were generated, and the volume of old and new bone calculated. For density calculations, DICOM files were imported into ImageJ (NIH, Bethesda, MD) and mean Hounsfield Units (HU) were obtained from a cylindrical volume of 0.045 cm^3^ consisting of the defect and 0.03 cm^3^ outside of the defect in the native bone as an internal control. The ratio of mean density of defect/mean density of native bone was then calculated for each condition.

In vivo rabbit cranium imaging was performed at the University of California, Los Angeles (UCLA) Crump Preclinical Imaging Technology Center. During the survival period at 3, 6, and 9 months after implantation, rabbits were injected with 1-5 mg/kg alfaxalone before anesthesia was maintained with 3-5% isoflurane gas. Each rabbit was placed on the imaging bed of a GNEXT Positron Emission Tomography and Computed Tomography (PET/CT) scanner (Sofie Biosciences, Culver City, CA). The rabbit cranium was scanned with a source voltage of 80 kVp, Intensity (µA) of 150, using 720 projections, and a 1-minute scan time. Three-dimensional (3D) image reconstruction was performed using a Modified Feldkamp Algorithm and analyzed with Dragonfly software (Dragonfly 4.0, Object Research Systems Inc., Montreal,

Canada). 3D reconstructions were cropped and a 0.92 cm^3^ cylindrical region of interest (ROI) was placed on the defect. The Otsu segmentation method was used to distinguish bone from non-bone. The regenerated bone mass was subsequently calculated by multiplying the bone volume (mm^3^) and density (Hounsfield Units, HU) (21,22).

### Histology and immunohistochemistry

Skulls were removed 3 months or 9 months after the implantation of scaffolds, fixed in 10% formalin, decalcified, embedded in paraffin, and sectioned at 4 microns using standard techniques. The sections were deparaffinized and stained with hematoxylin and eosin. Images were captured with the Zeiss Axio Observer 3 inverted microscope with the ZEN 2.3 Pro software (Zeiss, Oberkochen, Germany) and analyzed qualitatively.

### Reference point indentation for in vivo rabbit cranial defects

Explanted rabbit skulls at 3 and 9 months were fixed and tested with the BioDent reference point indentation device (Active Life Scientific, Santa Barbara, CA) according to manufacturer’s instructions. Indentations were conducted in 10 areas of native bone and 20 areas of regenerated bone at a force of 2N, an indentation frequency of 2 Hz, and 10 indentation cycles at a touchdown force of 0.1 N using a probe assembly type BP2. Indentation data was analyzed with the BioDent software for the total indentation distance (TID), first cycle indentation distance (ID1st), loading slope (LS), and unloading slopes (US). Toughness, or resistance to fracture, was determined by the total distance of indentation (TID) reached by the test probe and the first cycle indentation distance (ID1st). Relative stiffness was determined by the loading (LS) and unloading (US) slopes of the force (N) to displacement (µm) curves. To minimize differences in the thickness of bone for each animal as well as the bone healing capabilities, data from each cranial defect was internally controlled with the native calvarial bone.

### Scanning Electron Microscopy (SEM) imaging

Morphological characterization of was visualized using Scanning Electron Microscopy (SEM) (Supra 40VP SEM, ZEISS, Germany). Explanted rabbit skulls at 3 months and 9 months were fixed in 10% formalin overnight. Next, samples were rinsed with PBS, decalcified using a 0.5 M ethylenediaminetetraacetic acetic (EDTA) disodium salt solution, and stored in 70% ethanol. Subsequently, samples were placed inside a critical-point dryer (AutoSamdri 810 Critical Point Dryer, Tousimis, USA). Dried samples were then coated with a gold target (Ted Pella, USA) using a sputter coater (Pelco SC-7, Ted Pella, USA) and imaged.

### Statistical Analysis

Differences between *ex vivo* microCT and reference point indentation analyses were compared with a Kruskal-Wallis test due to non-normal distributions of the data. Pairwise comparisons were performed using the Dunn’s test with a Bonferroni adjustment. Mean differences between *in vivo* microCT analyses were compared with a one-way ANOVA with posthoc comparisons under the Tukey criterion. A p-value less than 0.05 was considered significant. All statistical analyses were performed using SPSS software Version 28 (SPSS, Inc., Chicago, IL).

## Results

We utilized a critical-sized rabbit cranial defect model to compare the differences in bone healing between unreconstructed defects (Defect only), defects reconstructed with PDLLA implants (Defect-PDLLA), defects reconstructed with MC-GAG scaffolds (MC-GAG), and defects reconstructed with MC-GAG and PDLLA implants (MC-GAG-PDLLA). Initial bone healing assessment was conducted on explanted skulls at 3 months. For a comprehensive understanding of long-term effects, *in vivo* microCT imaging was subsequently performed at 3, 6, and 9 months, with further examination of explanted skulls at 9 months.

### Mineralization, mechanical, and morphological properties of rabbit calvarial defects reconstructed with MC-GAG and PDLLA implants at 3 months

At 3 months, the PDLLA implants were not degraded. *Ex vivo* microCT scanning of the explanted skulls was conducted to analyze mineralized content within the defect. A qualitative improvement in mineralization was observed in skulls reconstructed with MC-GAG, both with and without PDLLA implants. Mineralized content was quantified using median gray values from the microCT scans within a fixed volume of the defect, internally corrected by the density within a fixed volume in the native calvarium for each animal (Defect/Native mineralization ratios). Comparisons of Defect/Native mineralization ratios revealed significant differences between groups [H(3)=332.20, p<0.001]. Both MC-GAG [median(IQR): 0.43(1.41)] and MC-GAG-PDLLA [median(IQR): 0.75(2.04)] exhibited significantly higher mineralization ratios compared to unreconstructed defects [median(IQR): 0.31(0.17), p<0.0001]. The addition of a PDLLA implant also led to significantly higher mineralization; unreconstructed defects with PDLLA implants [median(IQR): 0.76(0.36)] showed higher ratios than defects only (p<0.0001). Similarly, mineralization ratios were significantly higher in MC-GAG-PDLLA compared to MC-GAG alone (p<0.001).

To confirm the presence of mineralization, histologic analyses of explanted defects were performed at the interface between native bone and defect. In the unreconstructed defect, predominantly fibrous, non-mineralized soft tissue was observed. In contrast, defects reconstructed with MC-GAG exhibited a more intricate network of trabecular, mineralized content. MC-GAG-PDLLA reconstructions also demonstrated more regenerated bone compared to unreconstructed defects. The mineral content in MC-GAG-reconstructed defects appeared less structurally organized compared to MC-GAG-PDLLA. The latter displayed an organization closely resembling that of the surrounding native bone.

Next, we assessed the mechanical properties of the defect using reference point indentation with two indicators of strength, namely the initial cycle indentation distance (ID1st) and total indentation distance (TID), along with two measures of stiffness, represented by the average relative unloading slope (US) and loading slope (LS). To mitigate variances in bone healing and thickness across animals, all bioindentation measurements were normalized as relative ratios between the defect and native bone.

The total indentation distance (TID), which is inversely correlated to microfracture resistance, was found to be different among the groups [H(3)=69.80, p<0.001]. Both MC-GAG [median(IQR): 2.13(1.19)] and MC-GAG-PDLLA [median(IQR): 2.10(1.97)] demonstrated lower TID compared to defect only [median(IQR): 3.09(1.66), p<0.0001 and p=0.03, respectively] and defect-PDLLA controls [median(IQR): 4.63(0.78), p<0.0001]. The 1^st^ cycle indentation distance (ID1st), which is inversely correlated to mineralization and density, was also found to be significantly different among the groups [H(3)=69.58, p<0.001]. MC-GAG [median(IQR): 2.06(1.16)] and MC-GAG-PDLLA [median(IQR): 2.18(1.92)] exhibited significantly lower relative ID1st values compared to defect-PDLLA controls [median(IQR): 4.57(0.93)], and MC-GAG alone was also significantly lower than defect only controls [median(IQR): 2.90(1.53), p<0.0001]. No differences were found in relative TID and ID1st values between MC-GAG and MC-GAG-PDLLA (p=0.456 and p=0.125).

Average relative unloading slope (US) and loading slope (LS) were also found to be significantly different among the groups [H(3)=130.76, p<0.001; H(3)=123.63, p<0.001, respectively]. In both measurements, MC-GAG [US median(IQR): 0.34(0.39) and LS median(IQR): 0.34(0.46)], and MC-GAG-PDLLA [US median(IQR): 0.78(0.38) and LS median(IQR): 0.64(0.46)] were significantly higher than defect only [US median(IQR): 0.20(0.19) and LS median(IQR): 0.19(0.24)] and defect-PDLLA controls [US median(IQR): 0.09(0.03) and LS median(IQR): 0.09(0.03), p<0.001]. MC-GAG-PDLLA also demonstrated higher US and LS values compared to MC-GAG (p<0.001).

To evaluate the morphological characterization of native bone, defect only, defect-PDLLA, MC-GAG, and MC-GAG-PDLLA at 3 months, scanning electron microscopy (SEM) images were taken. The surface appearance of the regenerated bone was shown in Figure 3E. Notably, MC-GAG-PDLLA demonstrated a similar interconnected porous structure and composition compared to native bone, whereas this was not visible in the defect only, defect-PDLLA and MC-GAG groups.

In combination, the micro-CT, histologic, mechanical, and morphological analyses suggested that the combination of MC-GAG with PDLLA was more efficient at mineralization compared to MC-GAG scaffolds.

### Long-term outcomes of rabbit calvarial defects reconstructed with MC-GAG and PDLLA implants

To assess the long-term effects of MC-GAG-PDLLA on mineralization, longitudinal *in vivo* rabbit skull CT scanning was conducted at 3, 6, and 9 months. There was a significant difference in total Hu between groups [F(3,10)=8.07, p=0.005], with higher values in MC-GAG-PDLLA compared to in MC-GAG-PDLLA compared to both the defect only and defect-PDLLA controls.

*Ex vivo* microCT scanning of the explanted skulls was conducted to analyze mineralized content within the defect. A qualitative improvement in mineralization was observed in skulls reconstructed with MC-GAG, both with and without PDLLA implants. Mineralized content was quantified using median gray values from the microCT scans within a fixed volume of the defect, internally corrected by the density within a fixed volume in the native calvarium for each animal (Defect/Native mineralization ratios). Comparisons of Defect/Native mineralization ratios revealed significant differences between groups [H(3)=346.133, p<0.001]. Both MC-GAG and MC-GAG-PDLLA exhibited significantly higher mineralization ratios compared to unreconstructed defects. The addition of a PDLLA implant also led to significantly higher mineralization; unreconstructed defects with PDLLA implants showed higher ratios than defects only (p<0.001). Similarly, mineralization ratios were significantly higher in MC-GAG-PDLLA compared to MC-GAG alone (p<0.001).

To assess mineralization, histological evaluation was conducted at the junction between native bone and the defect site. In areas without reconstruction, the tissue observed was mainly fibrous and lacked mineralization. In contrast, defects treated with MC-GAG showed a more complex network of mineralized trabecular structures. Additionally, MC-GAG-PDLLA-treated defects exhibited greater bone regeneration than those left unreconstructed. However, the mineral structure in MC-GAG-only samples was less organized than that seen with MC-GAG-PDLLA, which more closely mirrored the architecture of native bone.

At 9 months, the explanted skulls showed full degradation of the PDLLA implants and were subsequently assessed for their mechanical properties. Both the TID and ID1st were found to be different among the groups [H(3)=38.972, p<0.001 and H(3)=34.907, p<0.001, respectively]. MC-GAG-PDLLA consistently exhibited lower relative values for these measurements compared to all other groups.

Similarly, the stiffness measurements, US and LS, demonstrated significant differences between the groups [H(3)=54.531, p<0.001 and H(3)=53.952, p<0.001, respectively], with higher values in MC-GAG-PDLLA compared to all other groups.

Taken together, these data suggested that MC-GAG-PDLLA promoted enhanced mineralization at 6 months and exhibited improved mechanical properties at 9 months compared to all other groups.

## Discussion

Rabbit cranial defects were induced and reconstructed with an MC-GAG scaffolds stabilized by a Poly(DL-lactide) (PDLLA) plate onlay fixed with PDLLA screws, and were compared to empty defects without treatment, defects reconstructed with a press-fitted MC-GAG scaffold alone, and unreconstructed defects treated with the PDLLA onlay implant alone. The explanted skulls were assessed at 3 and 9 months following implantation, with *in vivo* microCT analyses conducted at 3, 6, and 9 months. At 3 months, the explanted skulls revealed the MC-GAG scaffold promoted mineralization in comparison to the PDLLA-treated and empty defects, with even greater mineralization observed for the MC-GAG-PDLLA combination scaffold. Histological analysis of the interface between the reconstruction and native bone revealed that MC-GAG promoted the development of an intricate network of trabecular mineralized content compared to the untreated defect—primarily featuring fibrous, non-mineralized tissue—with enhanced structural organization more closely representative of nearby native bone for MC-GAG-PDLLA. Examining the biomechanical properties of the treated skulls revealed enhanced hardness and microfracture resistance of both MC-GAG alone and MC-GAG-PDLLA compared to the untreated and PDLLA-treated defects, with significantly greater measures of stiffness for MC-GAG-PDLLA. While no difference in regenerated bone mass measured by *in vivo* CT was detected between the treatment groups after 3 months, MC-GAG-PDLLA featured the greatest bone mass at 6 months, and both MC-GAG-containing scaffolds demonstrated enhanced bone regeneration compared to the PDLLA and empty defects at 9 months. While the PDLLA plates were still discernible at the 3 month mark, they were completely resorbed by 9 months. Analyses of the mechanical properties of explanted skulls after 9 months revealed that both strength and stiffness were greatest and most similar to native bone for MC-GAG-PDLLA. Together, these results demonstrate that the reconstruction of a bone defect with a MC-GAG scaffold fixated with PDLLA significantly enhanced long-term mineralization and bone regeneration compared to a scaffold consisting of either MC-GAG or PDLLA alone.

In the use of bone-grafts or bone-substitution materials to reconstruct bone defects and fractures, the interface between native bone and the scaffold material dictates much of the bone-healing and regeneration process; to ensure the stability of this interface, prior efforts have incorporated fixation components, and have demonstrated their significance in promoting scaffold resorption and subsequent bone deposition (23–31). Increased gap size and micromotion at the interface mitigates bone regeneration, and bulk scaffold instability at the defect site may lead to partial or complete detachment resulting in bone step offs and irregular healing (32,33). In addition to promoting union, rigid fixation reduces excessive strain to the defect site that would otherwise hinder bone regeneration (34), and may also provide functional benefits to patients through quickening return to activity and expediting the healing process by holding the bone replacement material in place, and protecting the defect site or deeper structures exposed by the defect. Similarly to the results described here, Kwon et al. also achieved enhanced bone regeneration through the addition of fixation mechanisms. The authors reconstructed critical sized rat calvarium defects with a polydopamine-laced hydroxyapatite collagen calcium silicate (HCCS-PDA) nanocomposite scaffold and compared the results of unfixed defect sites to those fixated with a titanium mesh onlay to prevent axial movement, and Gelfoam at the interface to prevent radial movement. After 12 weeks, osteointegration was significantly higher for the fixation-reinforced defects, which also demonstrated greater resistance to push-out forces (35).

While fixation broadly promotes bone healing, the specific method of fixation can also differentially influence this process (36–38). For example, Ferguson et al. demonstrated that the position and profile of the fixation plate differentially influence the bone healing outcome in mandibular reconstruction (39). Fixators with mechanical properties that differ significantly from those of bone—for example, metal fixators such as titanium or steel plates and screws—may impede bone healing as a result of excess stress shielding (40). Stress shielding refers to the alleviation of typical stress experienced by bone at the site of treatment due to the use of fixation materials which differ in mechanical properties and stiffness compared to native bone (41). Because mechanical stimulation of a bone defect site influences the healing response, the approach to fixation must not be too rigid or flexible (42). In a critical segmental defect model in sheep, Pobloth et al. showed that the bone regeneration facilitated by defect reconstruction with a soft titanium-mesh scaffold and onlay fixation plate was reduced when the stiffness of the fixation plate was increased (43).

Efforts have shifted from static fixation to dynamic fixation, wherein the mechanical support conferred by the fixator is reduced overtime due to material degradation or replacement (44,45). The use of non-degradable fixation materials poses the risk of malposition and palpability of the hardware, and may require removal, increasing the risk for further injury and pain. Additionally, degradable fixation devices are suggested to enhance stress protection due to their similarity of elasticity compared to bone and provide gradual return of physiological stress to the healing bone over time (46–48). Chacon et al. compared the reconstruction of mandibular defects with rabbit tibial onlay bone grafts, with and without fixation with biodegradable screws (Lactosorb). Compared to the non-fixated control — which demonstrated no integration into the bone—the graft fixated with the screw demonstrated smooth integration at the graft-bone interface, with significantly greater thickness of the graft site (47).

Among biodegradable materials available for clinical implementation, PDLLA is frequently employed in bone fixation due to its strength, biocompatibility, and amenability to replacement over time with endogenous bone (49–52). The PDLLA- based fixator plate (ResorbX) utilized here was integrated using ultrasound-aided fixation (Sonic Weld Rx) of PDLLA pins (Sonic Pin Rx); this approach enhances fixation stability via simultaneous welding of the plate, screw, and bone, while also allowing for facile intraoperative handling and demonstrating no adverse reaction following implementation (53–55). Overall, PDLLA-based scaffolds and the SonicWeld system have achieved successful clinical implementation and are well tolerated in craniofacial procedures (48,56–59). Marco et al. utilized the same ultrasound-based system to similarly implement PDLLA onlay membranes for fixation and investigated their impact on bone regeneration within pig calvarial defects reconstructed with autogenous bone particulates derived during defect creation. The authors compared untreated (empty) defects, defects treated with PDLLA alone, bone particulates alone, and a combination of PDLLA and bone particulates. After 40 days of implantation, the empty and PDLLA-treated defects demonstrated incomplete bone formation and dense, fibrous connective tissue, while both bone particulate samples showed similarly complete bone formation (52). While the authors did not identify an improvement in bone regeneration with the combination of the PDLLA fixator and the bone-substitution material, the observation period was much shorter than ours and did not exceed 40 days.

The ideal features of a bone-replacement material include ease of handling, proper cell, and tissue ingrowth, bioresorbability, osteoconductivity, stability at the defect site, and adequate mechanical strength throughout the healing process. Clinical implementation of grafts with stable fixation may be difficult within large bone defects when using stiff allografts or ceramic-based bone substitutes, such as hydroxyapatite (24). Without fixation, bone replacement material must typically be press-fitted into the defect site. While this may not be experimentally challenging, as defect sites are artificially introduced with relatively simple dimensions—for example, a perfectly circular defect—the dimensions of defects encountered in the clinic such as those incurred through trauma are likely more complex. As such, it is prudent to design an adaptable scaffold that may be used off-the-shelf with minimal or no modification or contouring necessary for press-fitting into a site. However, while soft polymer-based materials such as hydrogels may provide ease of handling and conformability, they may lack sufficient mechanical properties and integrity alone to support long term bone regeneration. To achieve the aforementioned features while also addressing the design shortcomings, we propose a multi-component or composite structure combining the bone-regenerative MC-GAG scaffold capable of adapting and conforming to the bone defect, with stable and dynamic fixation provided by the PDLLA plate and its initially greater mechanical strength compared to the scaffold. Our group and others have shown that scaffolds composed of mineralized collagen such as MC-GAG may be better suited to repairing large cranial defects in comparison to traditional cranioplasty materials (5,7,8,11,12,16,60–63).

Though the reconstruction of cranial defects with MC-GAG alone may not provide adequate cerebral protection, this can be resolved with the addition of the PDLLA fixation plate, which is demonstrated here to also promote new bone deposition more closely reflective of the thickness, strength, and stiffness of surrounding native bone. In addition to enhancing stability and protection of the defect site, the use of a biodegradable fixation system as described here—allowing for initial stress protection, with gradual transferal of mechanical stress necessary for competent bone formation to the defect site—reduces concerns regarding stress shielding. While in this case the intervention described was implemented at a site with fairly minimal weight or load-bearing stress, the implementation of this system elsewhere—such as in long bones or the mandible—may require consideration of stress shielding.

The exact mechanism underlying the enhancement of bone regeneration via fixation at the bone defect site should be made the subject of further study. Possible explanations include enhanced interfacial stability and reduced micro- or macro- motion, temporally-regulated alleviation of mechanical strain at the defect site, protection of the healing defect from external forces, or some combination of these. Beyond its influence on mechanics and motion, the extent to which plate fixation improves vascularization and preserves blood supply to the bone site should be further investigated, as fixation stability has been shown to impact vascularization (64). It is possible that the enhanced bone regeneration and mineralization observed for the combination of MC-GAG and the PDLLA plate may result from intrinsic bone regeneration-promoting capacities of PDLLA—or its breakdown products—other than its use as a means of fixation. However, to the best of our knowledge, this has not been reported in the current literature. Additionally, as the specific fixation technique is known to influence clinical outcome, it may be worthwhile to explore other means of providing stabilization (39). This may be especially prudent considering that use of additional materials for fixation—including the plate and the screws implemented here—and the associated increase in overall collective area of the scaffold-bone interface introduces increased risk for infection (65). Future studies may implement a fixation approach that achieves the necessary mechanical stability without significantly increasing the contact area.

---

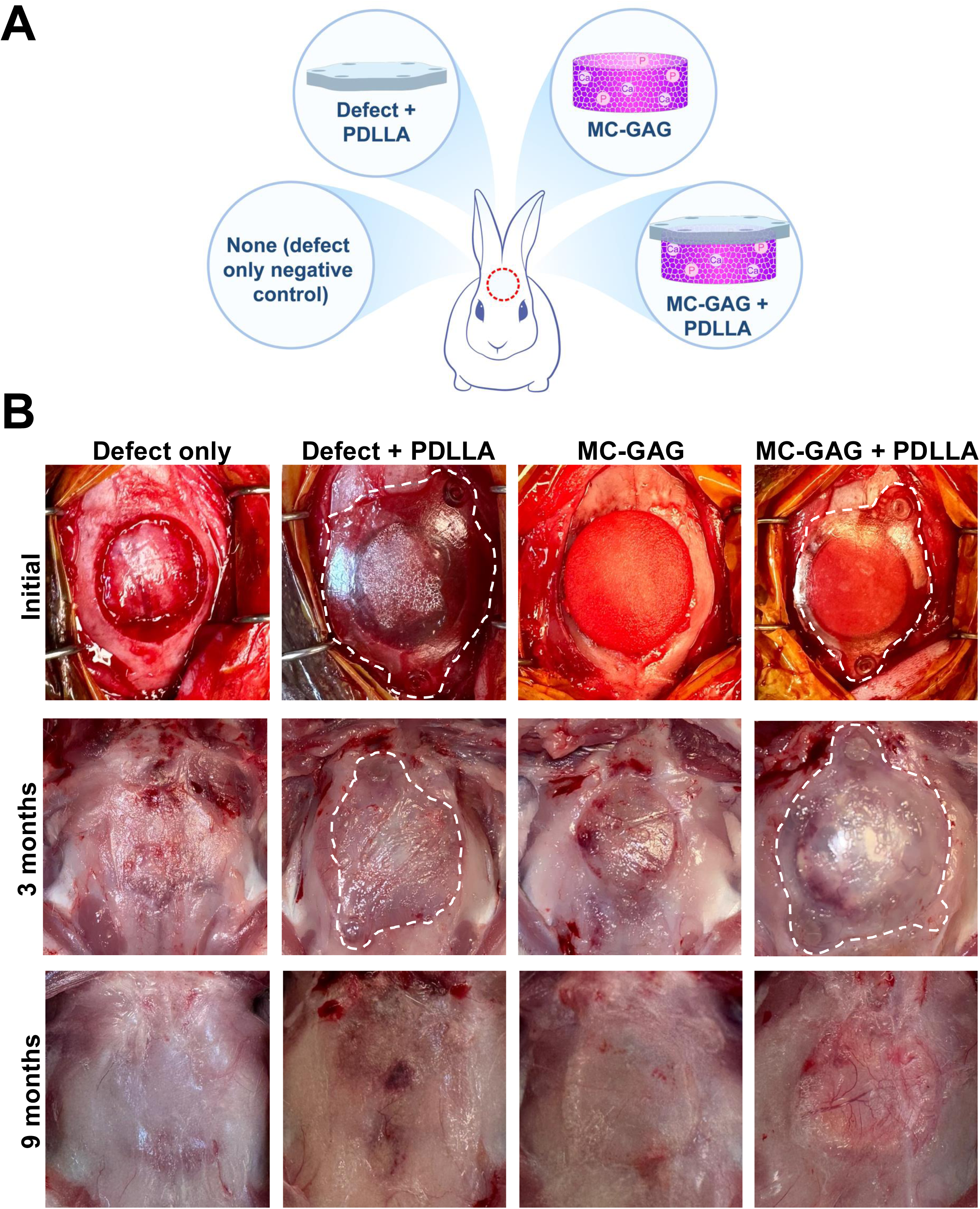

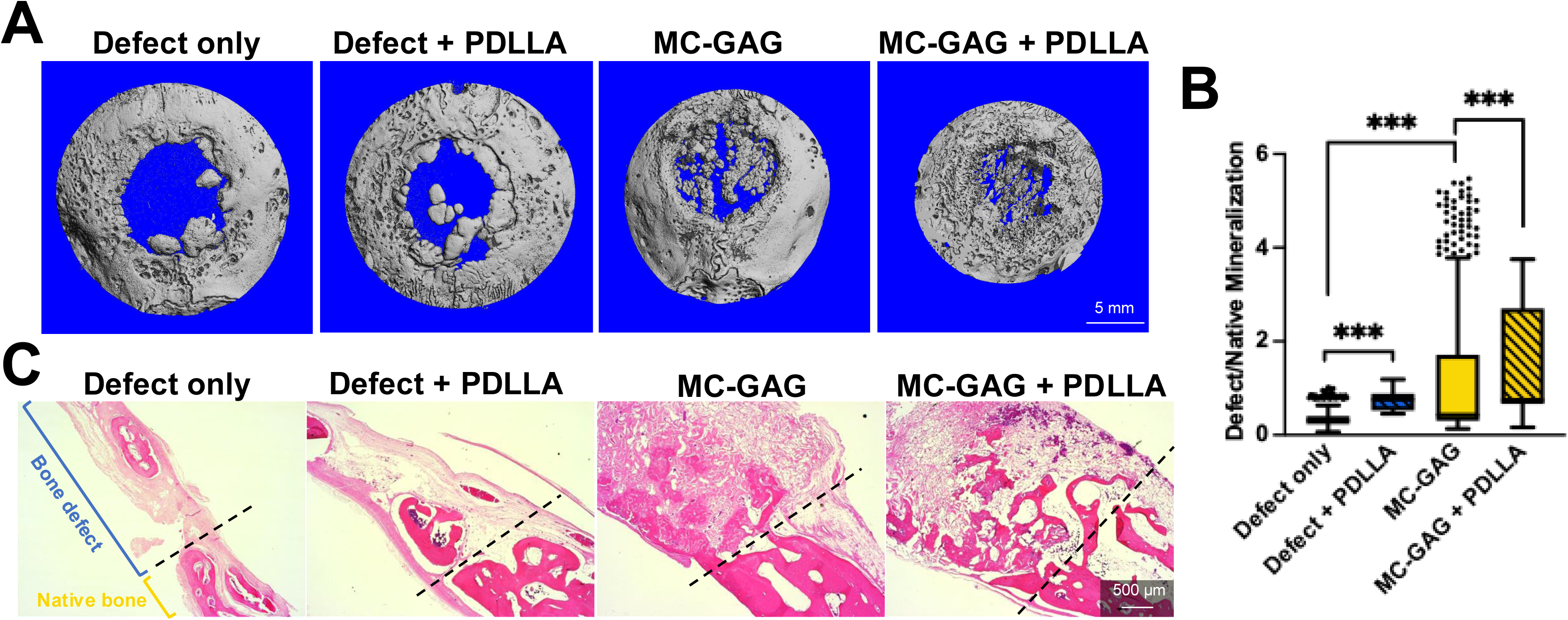

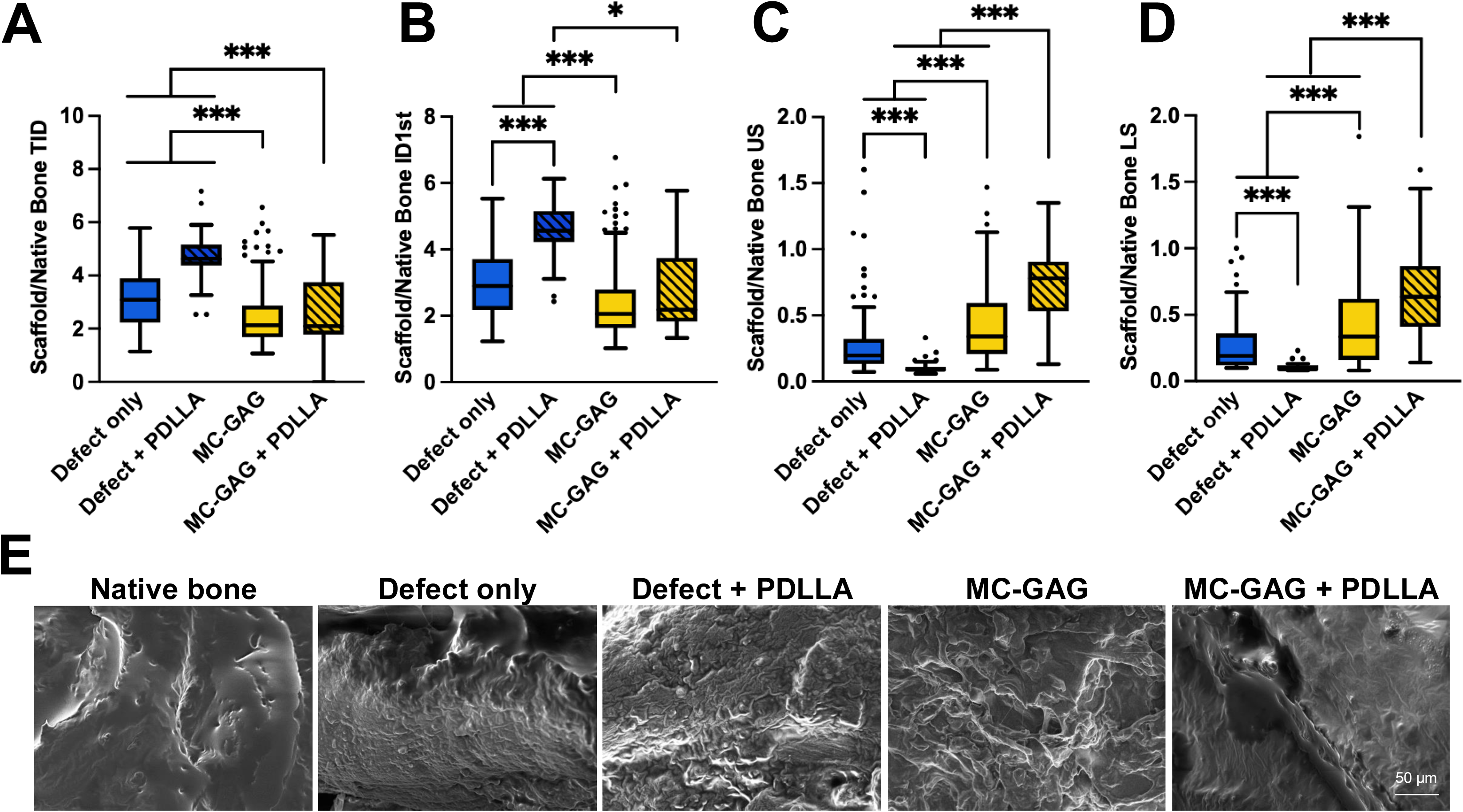

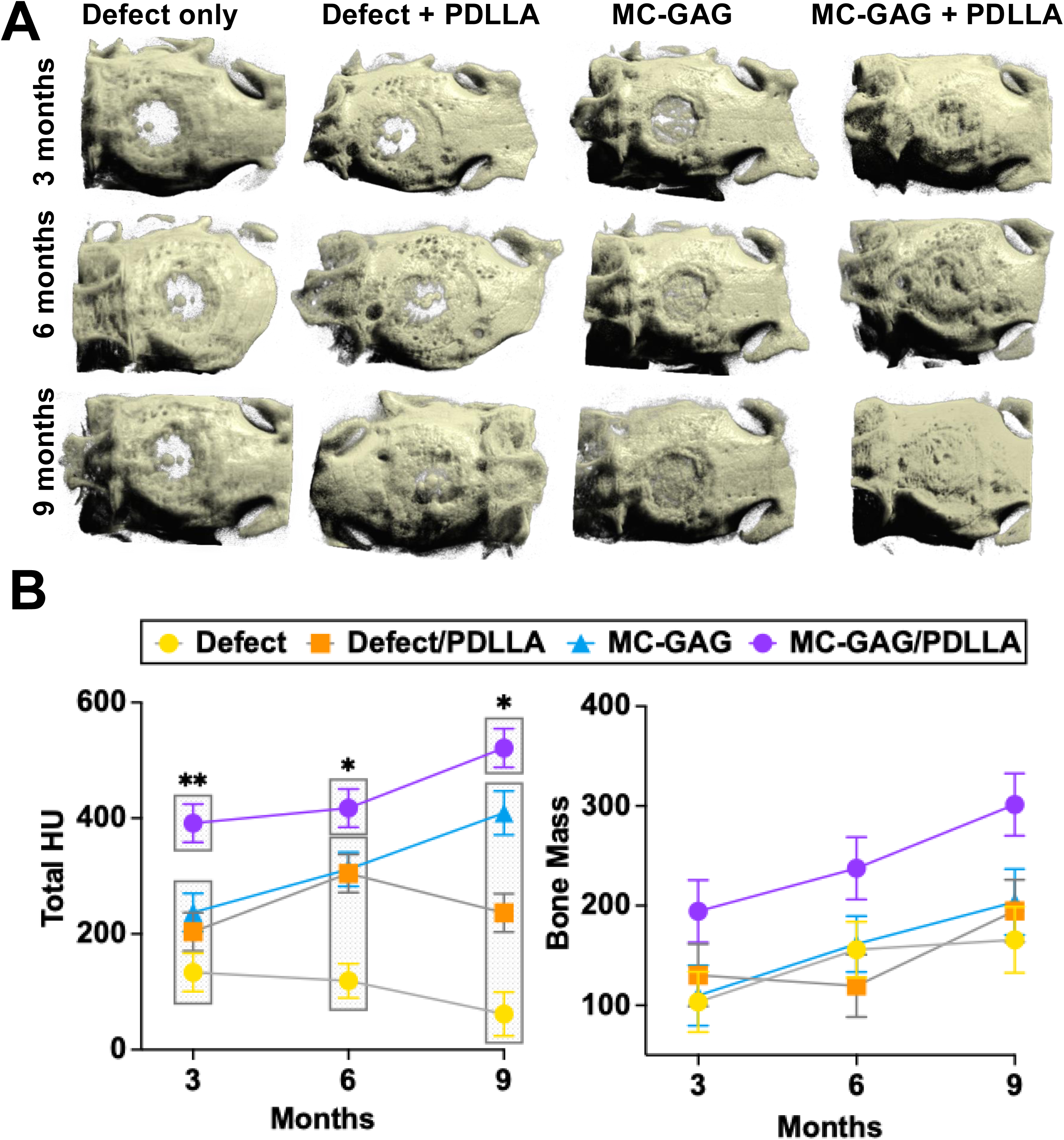

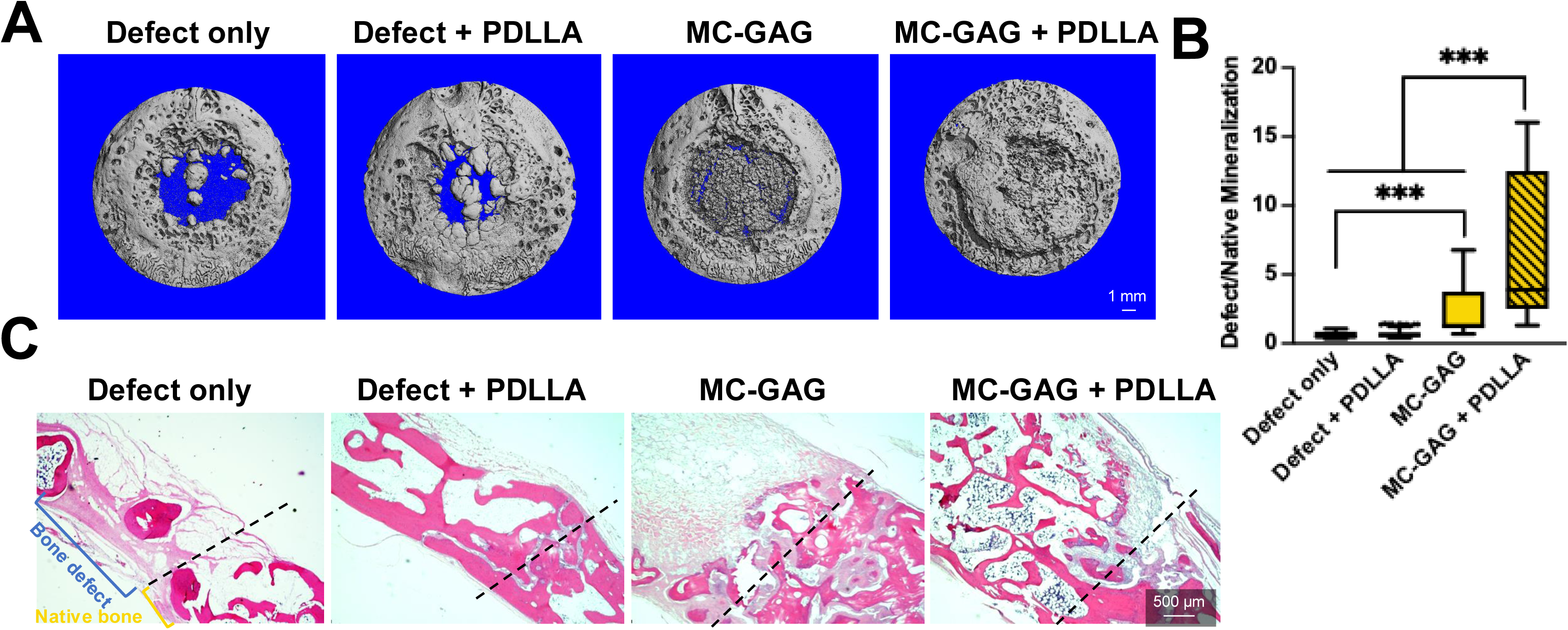

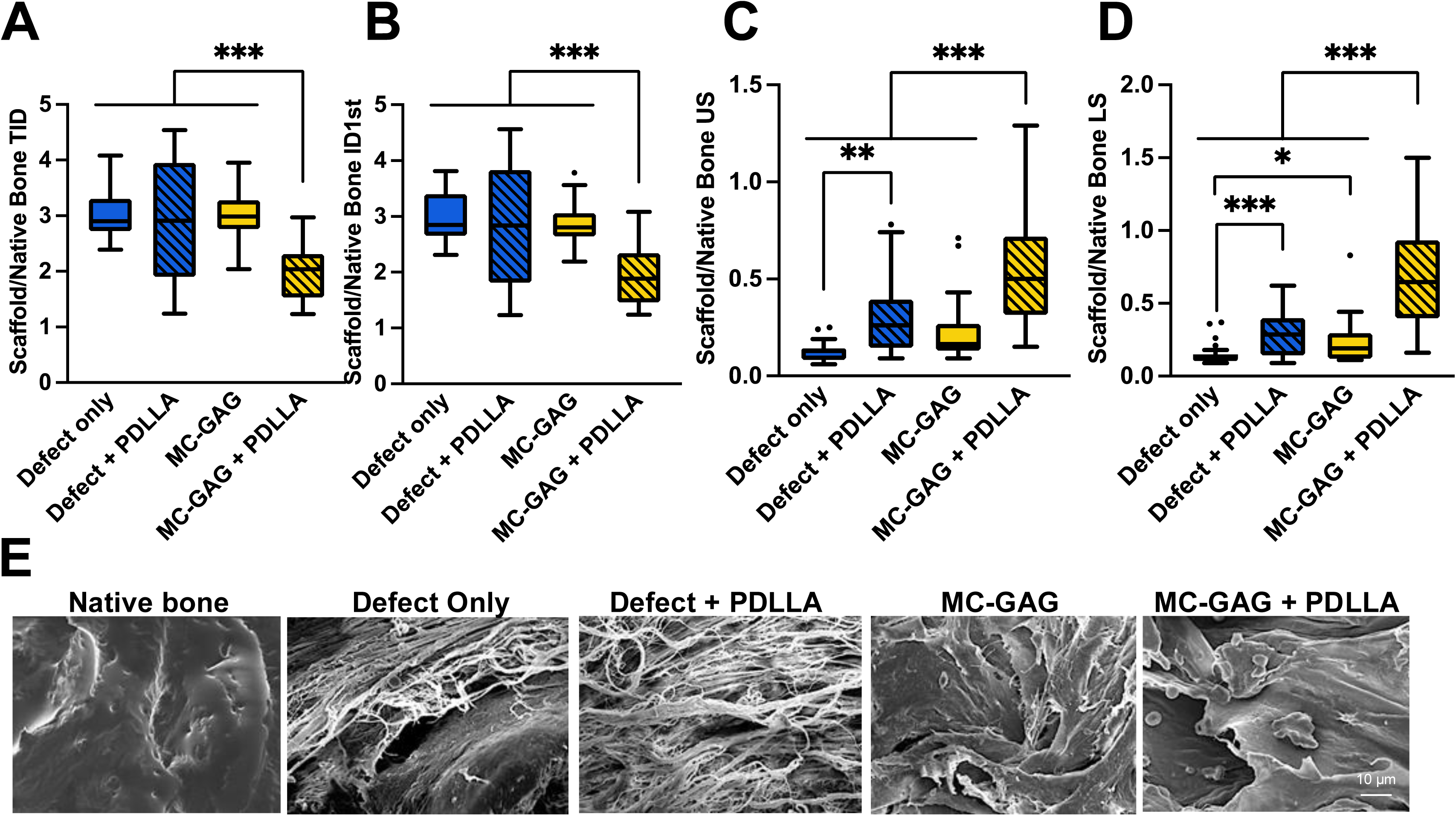

