## Supplementary material for "Resorbable Poly-(D,L)-Lactide Anchorage of Nanoparticulate Mineralized Collagen Materials Maximizes *In Vivo* Skull Regeneration": Figure Legends

**Figure 1. Critical-Sized Rabbit Cranial Defects Implanted with MC-GAG Scaffolds and Poly(DL-lactide) (PDLLA) Implants.** (**A**) Schematic representation of the experimental groups. Biparietal cranial defects (14 mm diameter) were surgically created using a hand-powered trephine and left unreconstructed (defect only), reconstructed with a PDLLA implant, MC-GAG scaffold, or MC-GAG scaffold with PDLLA implant. (**B**) Images depict the initial defect, reconstruction with a PDLLA implant, MC-GAG scaffold, and MC-GAG with PDLLA implant at the time of implantation (top), as well as at 3 (middle) and 9 months (bottom) post-implantation. At 3 and 9 months post-implantation, the rabbit skulls containing the respective implants were grossly examined and explanted. PDLLA implants (dashed line labeled) were completely resorbed by 9 months.

**Figure 2. In Vivo Rabbit Calvarial Regeneration in Cranial Defects Implanted with MC-GAG Scaffolds and Poly(DL-lactide) (PDLLA) Implants at 3 Months.** Representative images (**A**) and quantitative analysis (**B**) of *ex vivo* micro-CT scanning of explanted rabbit skulls 3 months after unreconstructed defects (Defect only) or defects reconstructed using PDLLA implants, MC-GAG scaffolds or MC-GAG with PDLLA. Quantitative data expressed as a ratio of mineralization within the defect corrected by mineralization within the surrounding native bone. Quantitative data analyzed using Krukal-Wallis tests with pairwise comparisons using Dunn’s test with Bonferroni adjustments. Representative images (**C**) of H&E staining of histologic sections within rabbit cranial defects explanted after 3 months. *, p<0.05; **, p<0.01; ***, p<0.001; ****, p<0.0001.

**Figure 3. Biomechanical Properties of Rabbit Cranial Defects Implanted with MC-GAG Scaffolds and Poly(DL-lactide) (PDLLA) Implants at 3 Months.**

Reference point indentation in explanted rabbit skulls with 14-mm biparietal calvarial defects unreconstructed (defect only), reconstructed with a PDLLA implant, MC-GAG scaffold, or MC-GAG scaffold with PDLLA implant at 3 months showing (**A**) first cycle indentation distance (ID1st), (**B**) total indentation distance (TID), (**C**) unloading slope (US), and (**D**) loading slope (LS). All measurements were expressed as a ratio of the defect to native bone to account for individual differences in biomechanical properties within specific animals. Representative scanning electron microscopy images (**E**) within rabbit cranial defects explanted after 3 months.*, p<0.05; **, p<0.01; ***, p<0.001; ****, p<0.0001.

**Figure 4. Long-Term In Vivo Rabbit Calvarial Regeneration Outcomes in Cranial Defects Implanted with MC-GAG Scaffolds and Poly(DL-lactide) (PDLLA) Implants.**

Representative images (**A**) and quantitative analysis (**B**) of *in vivo* micro-CT scanning of rabbit skulls 3, 6, and 9 months following unreconstructed defects (Defect only) or defects reconstructed using PDLLA implants, MC-GAG scaffolds or MC-GAG with PDLLA. Quantitative data are presented as bone mass, calculated as the product of bone density in total Hounsfield units (HU) and bone mass in mm^3^. Statistical analysis was performed using one-way ANOVA with posthoc comparisons under the Tukey criterion. *, p<0.05; **, p<0.01; ***, p<0.001; ****, p<0.0001.

**Figure 5. In Vivo Rabbit Calvarial Regeneration in Cranial Defects Implanted with MC-GAG Scaffolds and Poly(DL-lactide) (PDLLA) Implants at 9 Months.** Representative images (**A**) and quantitative analysis (**B**) of *ex vivo* micro-CT scanning of explanted rabbit skulls 9 months after unreconstructed defects (Defect only) or defects reconstructed using PDLLA implants, MC-GAG scaffolds or MC-GAG with PDLLA. Quantitative data expressed as a ratio of mineralization within the defect corrected by mineralization within the surrounding native bone. Quantitative data analyzed using Krukal-Wallis tests with pairwise comparisons using Dunn’s test with Bonferroni adjustments. Representative images (**C**) of H&E staining of histologic sections within rabbit cranial defects explanted after 3 months. *, p<0.05; **, p<0.01; ***, p<0.001; ****, p<0.0001.

**Figure 6. Biomechanical Properties of Rabbit Cranial Defects Implanted with MC-GAG Scaffolds and Poly(DL-lactide) (PDLLA) Implants at 9 Months.**

Reference point indentation in explanted rabbit skulls with 14-mm biparietal calvarial defects unreconstructed (defect only), reconstructed with a PDLLA implant, MC-GAG scaffold, or MC-GAG scaffold with PDLLA implant at 9 months showing (**A**) first cycle indentation distance (ID1st), (**B**) total indentation distance (TID), (**C**) unloading slope (US), and (**D**) loading slope (LS). All measurements were expressed as a ratio of the defect to native bone to account for individual differences in biomechanical properties within specific animals. *, p<0.05; **, p<0.01; ***, p<0.001; ****, p<0.0001.
